# The application of metagenomics in the detection of arboviruses in mosquitoes (Diptera: Culicidade). A systematic review

**DOI:** 10.1101/2025.01.28.635234

**Authors:** Everson dos Santos David, Shirley Vasconcelos Komninakis, Erique da Costa Fonseca, Anne Caroline da Silva Soledade, Karen Carmo dos Santos, Raimundo Nonato Picanço Souto

## Abstract

Advances in deforestation and climate change directly imply changes in habits and the distribution of Culicidae across the globe, especially mosquitoes of medical importance and the main vectors of arboviruses. The viral metagenomics technique can be an important tool in characterizing the viral diversity present in mosquitoes. Thus, the aim was to identify evidence of the effectiveness of the viral metagenomics technique in detecting arboviruses in mosquitoes. This is a systematic review based on the PRISMA 2020 protocol. The research was carried out in five electronic databases: LILACS, PubMed, SciELO, Scopus and Web of Science, chosen to include studies published in health and interdisciplinary areas, as well as a complementary research on Google Scholar. Studies that used the viral metagenomics approach for the genomic evaluation of arboviruses found in mosquito samples were included, where the results demonstrated the presence of viral diversity and the identification of the genome of probable pathogenic viruses. The protocol was registered on the PROSPERO platform under the number CRD42024484713. 238 studies published in recent years were identified in the electronic databases. According to the inclusion/exclusion criteria, only 20 studies met the objectives for the systematic review. In all the studies, the viral metagenomics technique of genomic sequencing was applied to detect viruses, mainly those related to specific insect viruses, arboviruses, pathogenic viruses, animal viruses and plant viruses belonging to various viral families. Despite the challenges to be overcome in terms of the absence of reference sequences in genomic databases, the effectiveness of the metagenomics technique in characterizing the mosquito virome is clear from the studies, which broadens the understanding of viral diversity.

## INTRODUCTION

The spread of arboviruses around the world is favored by the high diversity of vectors, mainly mosquito species from the Culicidae family, due to their ability to transmit pathogens to humans and animals (1). Environmental, climatic and seasonal changes favor the occurrence and maintenance of transmission cycles of various arboviruses, occurring mainly through zoonotic routes that are maintained in enzootic transmission cycles, a predominant factor, which highlights the importance of public health (2).

In this way, it should be noted that arboviruses are widely distributed across the planet and are adapted to remote areas with high temperatures and low temperatures (3,4). There are approximately 600 species of arboviruses, of which approximately 150 are responsible for pathogens in humans and animals, considered accidental hosts, and the risk of infection is directly associated with exposure to environments where the viruses circulate (4–7).

Among the most relevant viruses are Dengue virus (DENV), Chikungunya virus (CHIKV), and Zika virus (ZIKV) transmitted mainly by the *Aedes aegypti* species (8); Yellow Fever Virus (YFV) transmitted in the wild by mosquitoes of the genus Haemagogus and Sabethes, and in the urban form by *Aedes aegypti* and *Aedes Albopictus*; Mayaro Virus (MAYV) transmitted by the *Haemagogus janthinomys* mosquito (9); The Oropouche virus (OROV) has already been found naturally infecting the *Coquillettidia venezuelensis* species (10); the Saint-Louis Encephalitis virus (SLEV) transmitted by mosquitoes of the Culex genus (5); and the Rocio virus (ROCV) isolated from the *Psorophora ferox* species (11).

In this sense, the viral metagenomics technique applied to different samples, such as plasma (6), feces (12), and in the various extreme habitats (13) has proven to be efficient in detecting viruses. Making its application in mosquito samples feasible, it can be effective in identifying arboviruses, due to the sensitivity of the technique in detecting the viral genomes that coexist in a given sample, contributing directly to the epidemiological surveillance of the main arboviruses (14).

In this sense, this important systematic review aims to seek evidence of the effectiveness of the viral metagenomics technique in detecting arboviruses in mosquitoes, enabling studies of the viral genetic variability of samples from different areas and the identification of emerging and re-emerging viruses capable of causing epidemics and even future pandemics.

## METHODS

This article is a systematic review based on the Preferred Reporting Items for Systematic Reviews and Meta-Analyses (PRISMA) 2020 protocol (15), adapted according to recent systematic review studies (16–18). It is registered in the International Prospective Register of Systematic Reviews (PROSPERO) under number CRD42024484713.

The formulation of the research question was developed using the acronym PICo as an auxiliary tool (19). PICo is made up of three components, namely Population or Problem, Interest and Context. Three aspects were addressed in this review based on these three components: detection of arboviruses in mosquitoes (problem), analysis of the effectiveness of the metagenomics technique (interest) and detection of arboviruses detected by the technique (context), elaborating the following research question: What is the evidence of the effectiveness of the viral metagenomics technique in detecting arboviruses in mosquitoes?

The research took place between April and July 2024, in five electronic databases: LILACS, PubMed, SciELO, Scopus and Web of Science, chosen to include studies published in health and interdisciplinary areas in English, Spanish and Portuguese, with no restrictions on the date of publication. In order to define the search terms, the selection of Health Sciences Descriptors (DeCS) and Medical Subject Headings (MeSH) in English was used for expansion, through the title and abstract.

The search strategy was based on the descriptors mentioned in figure 1. A manual search was also carried out on the references of the selected articles, as well as a Google Scholar search to identify relevant non-indexed studies.

**FIGURE 1:**
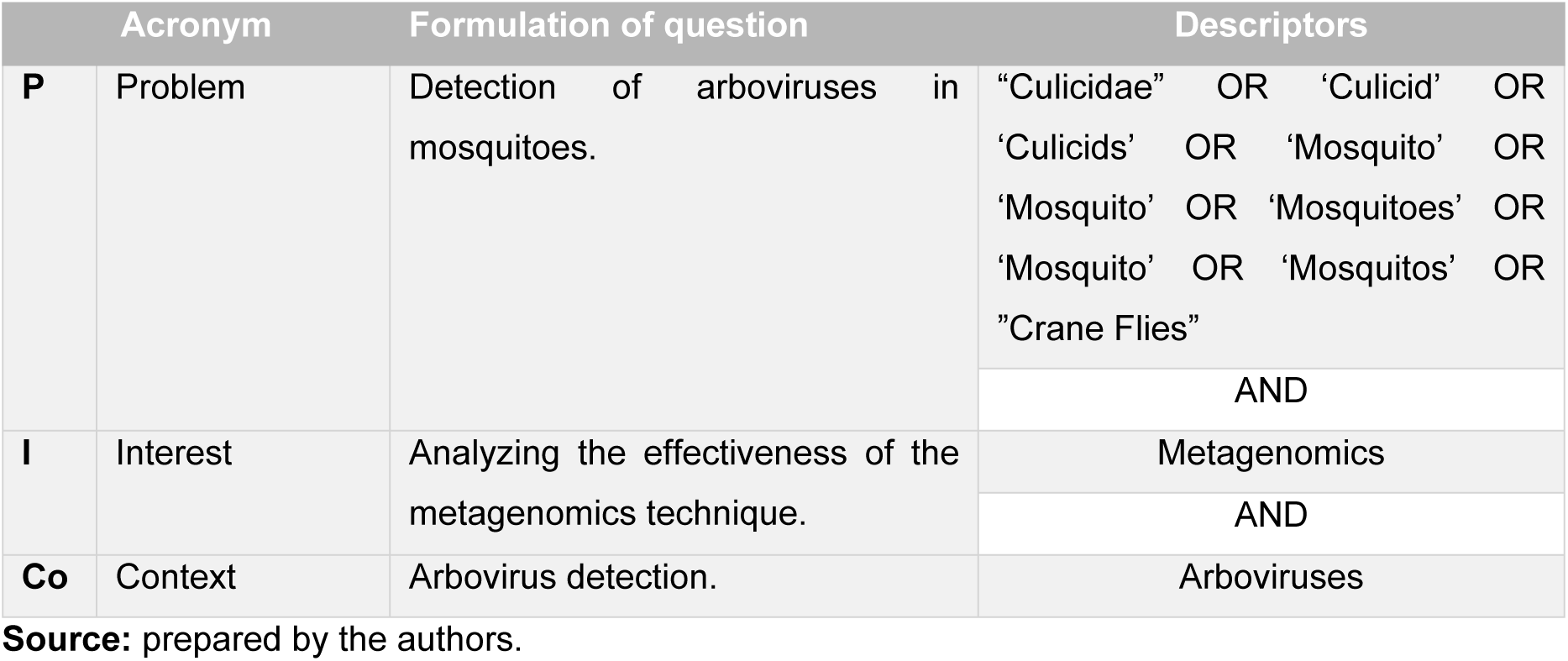
Formulation of the question based on the acronym PICo and the search strategy used for the systematic review.

### Eligibility criteria

Studies aimed at detecting arboviruses in mosquitoes were considered, specifically those belonging to the *Culicidae* family, especially those of genera considered to be of medical importance, such as *Anopheles*, *Culex* and *Aede*s, vectors of pathogens in humans. Studies that used the viral metagenomics approach for the genomic evaluation of all viruses existing in mosquito samples were included, where the results demonstrated the presence of viral diversity and the identification of the genome of probable pathological viruses.

Primary studies with an experimental design were included, whether prospective, retrospective or cross-sectional, carried out in any region of the world. Studies related to study objects such as ticks, blood samples or other insects analyzed by metagenomics were excluded.

### Data extraction and evidence synthesis

The files resulting from the database search were transferred to the Mendeley reference manager, version 1.18, in order to remove duplicates. The Rayyan QCRI platform (https://rayyan.qcri.org) was then used to select the articles. The first stage consisted of a title and abstract analysis carried out independently by two researchers (ESD and ECF). Disagreements were resolved independently by a third researcher (ACSS). After this stage, the full text was read independently by two authors (ESD and ECF). In addition, the reference list of the included studies was consulted to check for potentially eligible studies not identified in the database search.

Data extraction was performed independently by two reviewers (ESD and ECF) using an extraction form in Microsoft Excel 2016 (Microsoft, Redmond, WA, USA), with discrepancies resolved by consensus with a third researcher (ACSS). The data extraction form for the eligible studies included the following topics: genera of the *Culicidae* family, species, year of study, study objective, study area, *habitat* characteristics, country, mosquito distribution, number of pools, arboviruses identified in the samples, and the sequencer used in the research.

### Quality assessment

In order to assess the risk of bias, the SYRCLE (Systematic Review Center for Laboratory Animal Experimentation) tool for animal studies was used (20). This tool contains the following evaluation categories: selection bias, performance bias, detection bias, attrition bias, reporting bias and other sources of bias. Ten questions were applied to the articles included in the systematic review, the answers to which can be “YES”, indicating a low risk of bias, “NO”, indicating a high risk of bias, and “UNCERTAIN”, indicating an uncertain risk of bias. It is not recommended to calculate the sum score of each individual study using this tool.

## Results

### Characteristics of the included studies

According to the search strategies, 238 studies published in the electronic databases were identified, as shown in figure 2, following the criteria of the PRISMA protocol (15). After initially reading the titles and abstracts, 80 studies were screened for full text. According to the inclusion/exclusion criteria, only 17 studies were eligible for this systematic review.

**FIGURE 2:**
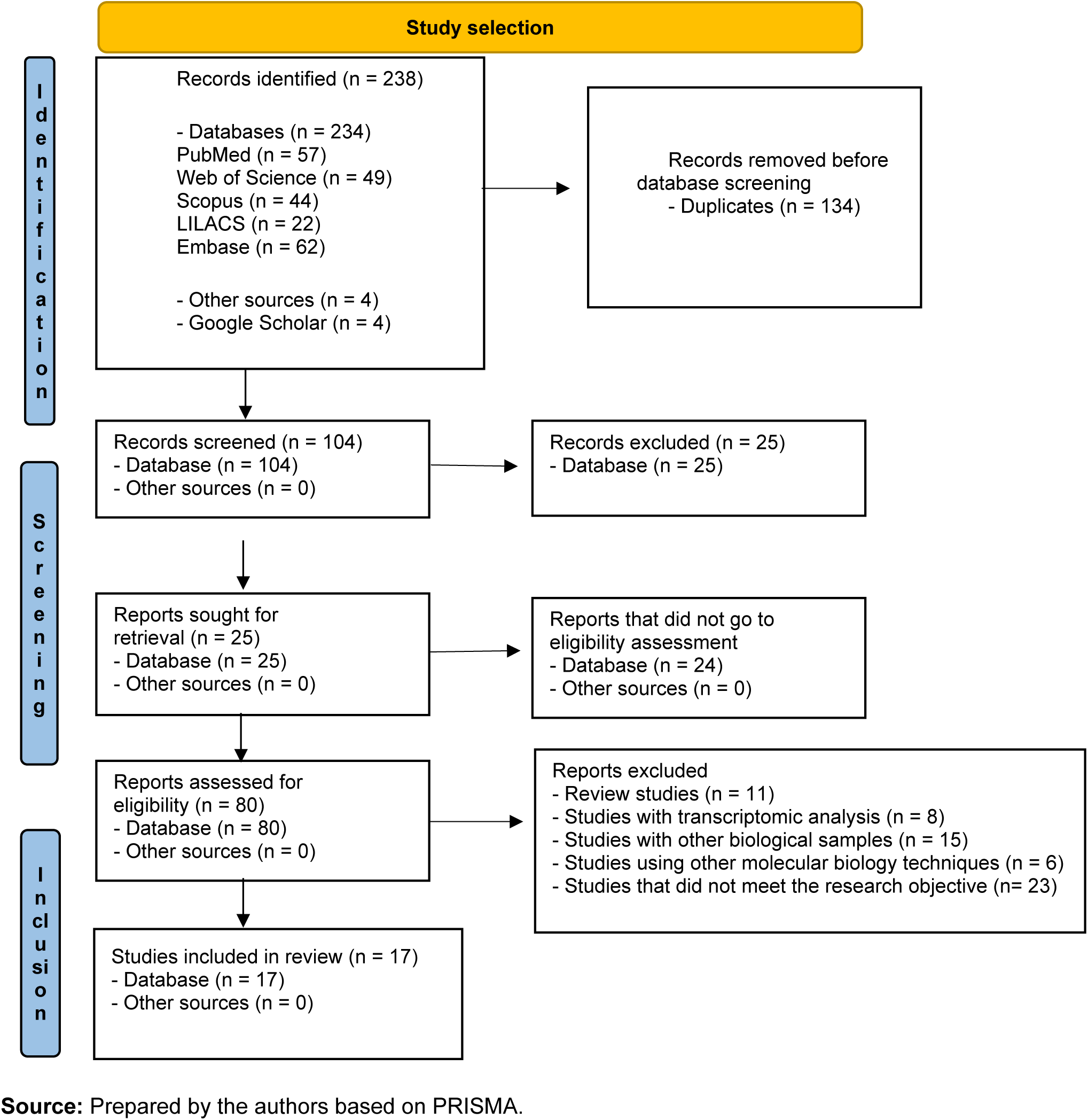
PRISMA 2020 flow diagram.

The main characteristics of the 17 studies included in this systematic review are shown in Table 1. The studies were published between 2016 and 2023, from different parts of the world, spread across five continents, and aimed to detect arboviruses in mosquitoes (Diptera: Culicidae) using the metagenomic approach.

**Table 1:**
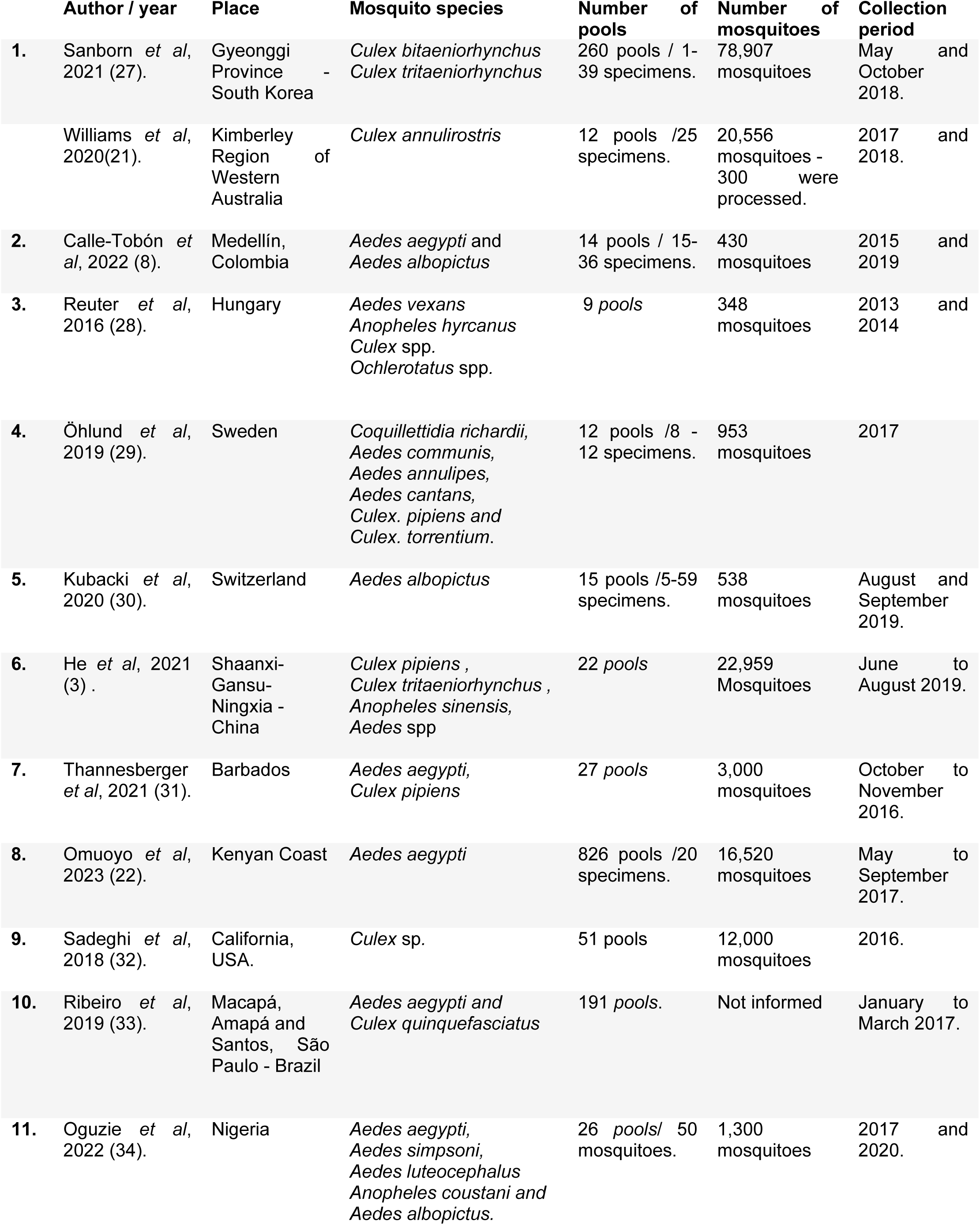
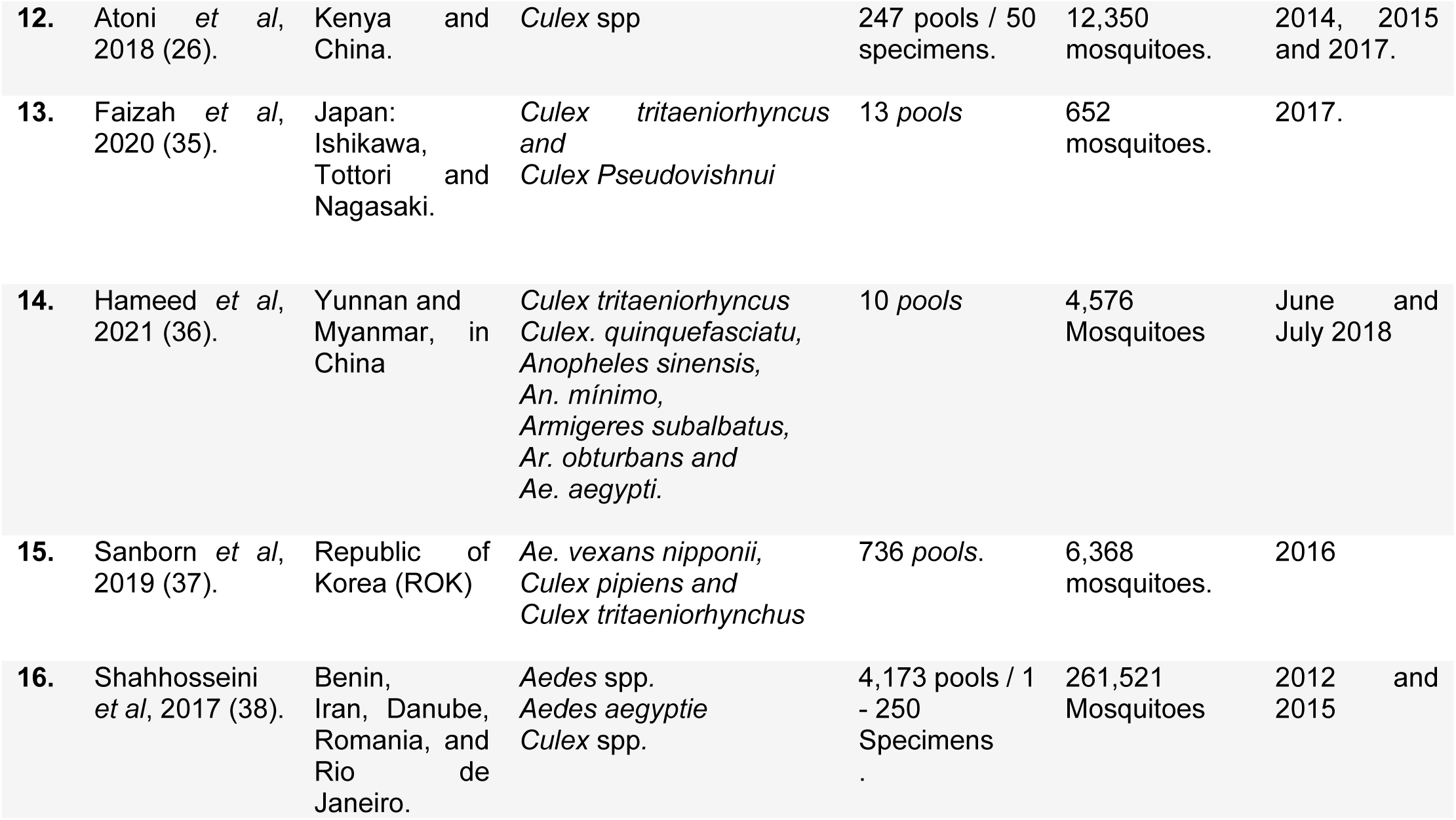
Main characteristics of the studies included in the systematic review.

However, the search strategies in the databases enabled the analysis of a range of results from the metagenomic approach, allowing not only the detection of arboviruses, but also knowledge of the diversity of viruses present in mosquitoes. Thus, it was demonstrated that the metagenomic technique has the potential to detect human and animal pathogens, without the need for cell culture or the use of animals, and viruses that are only found in mosquitoes (ISVs) (21).

The mosquito species shown in table 1 refer to the predominance of species in a given geographical area with an incidence of important and emerging arboviruses, areas chosen for their endemic importance. All the studies aimed to collect mosquitoes in areas with recorded arbovirus outbreaks.

The interest of the studies was the capture of female mosquitoes, due to the fact that they perform hematophagy, the act of sucking blood in search of nutrients necessary for the viability of reproduction. In this way, they transmit pathogens that are inoculated in humans and animals, causing diseases (22).

Mosquitoes are taxonomically classified in the order Diptera, suborder Nematocera and family Culicidae, with approximately 3,618 described species grouped into two subfamilies: Anophelinae and Culicinae. Thus, the eligible studies targeted two subfamilies, specifically the species belonging to the genus Anopheles, Aedes and Culex, because these vectors have the ability to transmit pathogens to humans and vertebrate animals, and are more efficient at transmitting pathogens (23,24).

Of the 17 eligible studies, 12 obtained results on the diversity of *Aedes mosquito viruses*, eight of which applied the metagenomics method to the Aedes aegypti species and two to the *Aedes albopictus* species. Both species are considered cosmopolitan due to their occurrence on all continents, where they thrive in the same types of breeding grounds, in urban environments (25).

Species of the genus Culex were predominant in 13 studies, as they are an efficient vector for transmitting West Nile virus, Rift Valley fever virus and Japanese encephalitis, as well as the parasitic disease bancroftian filariasis (26).

#### Viral Diversity

Of the 17 eligible studies, only seven showed positive results for the detection of human pathogens, such as Japanese encephalitis (JEV) (3,35,36,39), Dengue type 2 (26), Dengue type 3 (8), Zika virus (ZIKV) (31). Of the others, seven studies reported that they had not detected any human or animal pathogenic viruses in the samples. However, despite not detecting pathogens, the results showed the characterization of the viral diversity present in the different species of Culicidae, the focus of each study.

Three studies described the genetic characterization of new viruses, such as Reuter *et al.* (28) who identified three strains of a rhabdovirus isolated from three pools of mosquitoes of the genus Aedes and subgenus Ochlerotatus sp. collected in Hungary. Similarly, the studies by Shahhosseini *et al* (38) also focused on characterizing the genome and screening a new rhabdovirus, which was identified in *Ae. cantans* mosquitoes collected in Germany. And the studies by Ribeiro *et al.* (33) identified the occurrence of strains of the Hubei reo-like virus 7 (HRLV 7), known as an ISV (Insect-Specific Virus) detected in one pool of *A. aegypti* and five pools of *C. quinquefasciatus* in Brazil.

All the studies describe in their methodological procedures a flow that goes through the choice of collection environments (peri-domestic areas, forest areas, urban or rural regions, farms, corrals, etc.), mainly those with a history of pathogen incidence, mosquito capture, as well as storage and transport, taxonomic identification, pool organization, sample processing, genomic sequencing, bioinformatics analyses, and phylogenetic analysis. Few differences were observed in the methodological procedures, just the routine ones for each research laboratory.

The sample number of mosquitoes to compose the formation of the pools aims to elucidate the viromes of mosquitoes from a different location, so that it represents the viral diversity and abundance in a given period, and that it can reflect the compositions of the virome, to understand the dynamics of virus transmission in comparison to disease outbreaks (3).

In all the studies, the viral metagenomics technique of genomic sequencing was applied, which resulted in the detection of insect-specific viruses (ISVs), human pathogenic viruses, animal viruses, plant viruses, bacteriophages and environmental viruses with a small percentage, but which integrate the virome of several mosquito species (26). The various viral families are shown in the results column of table 2.

**Table 2:**
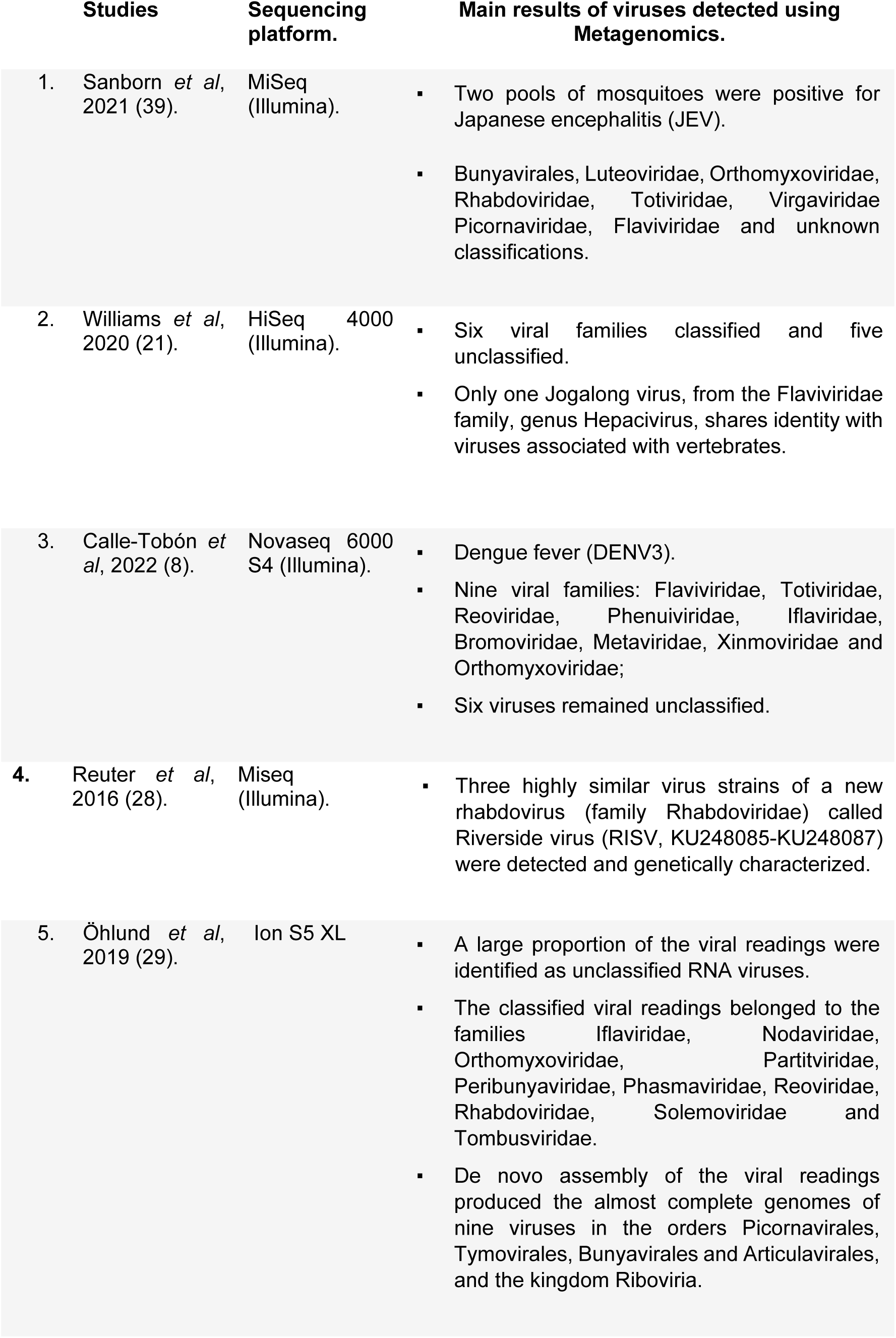
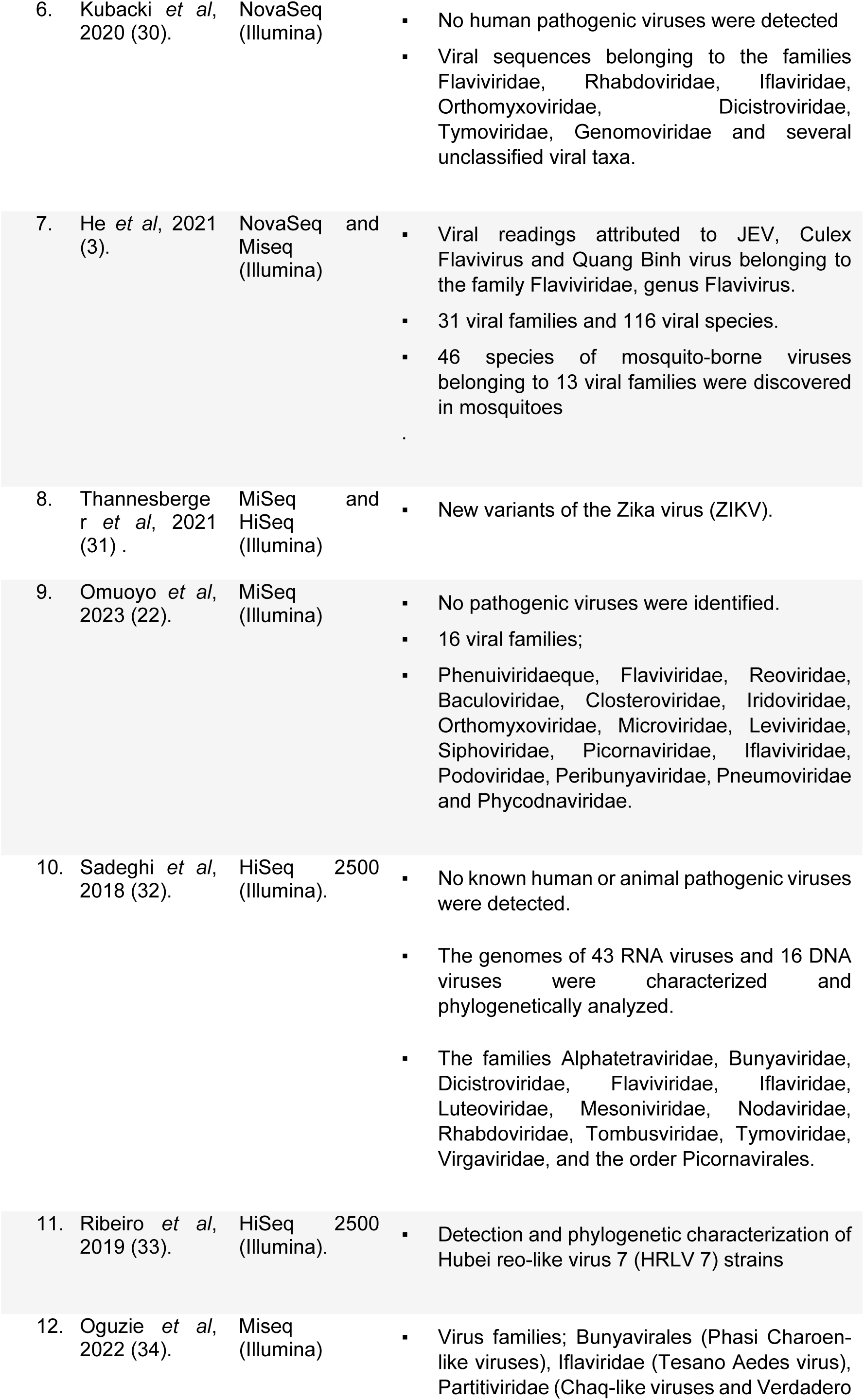
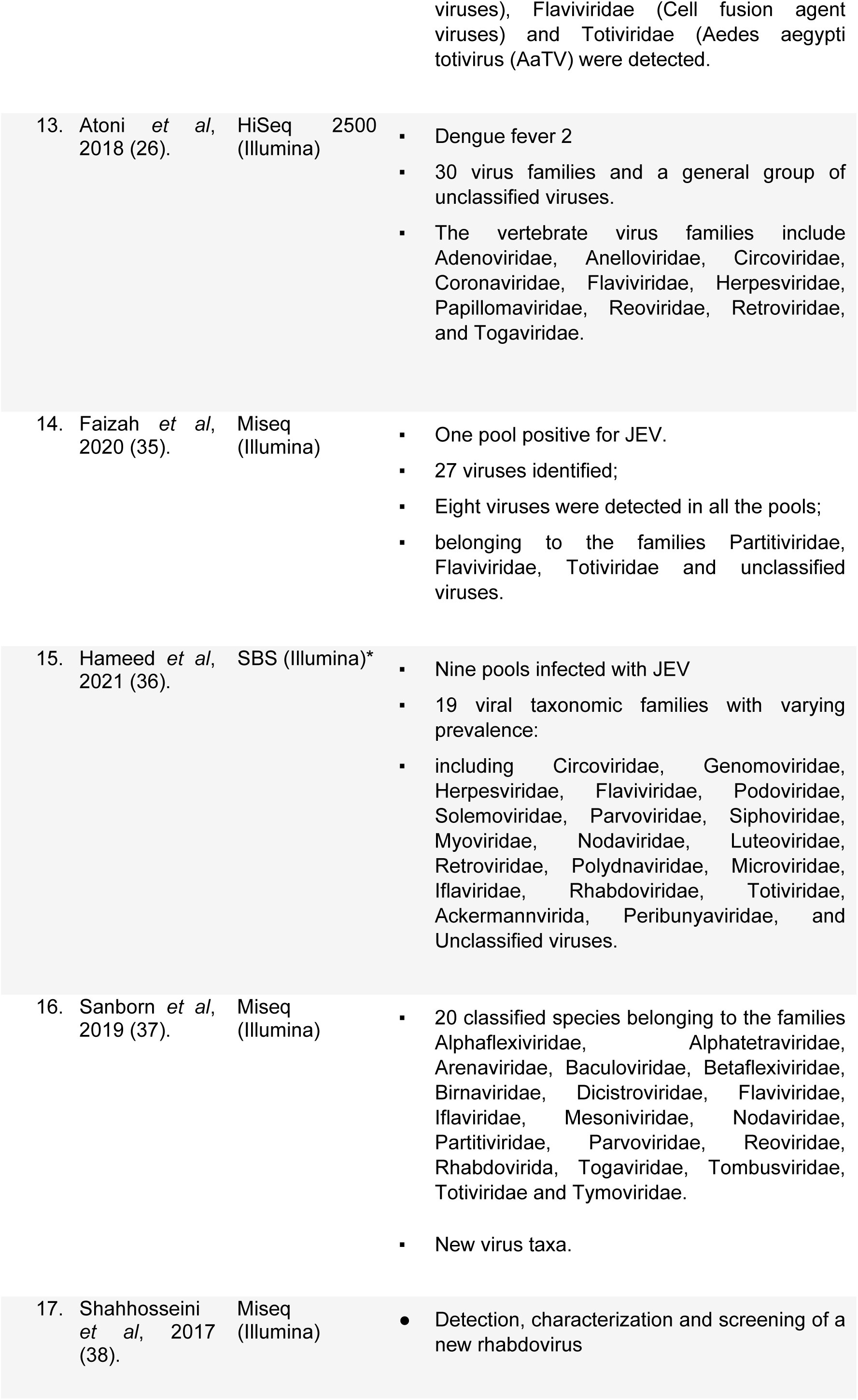
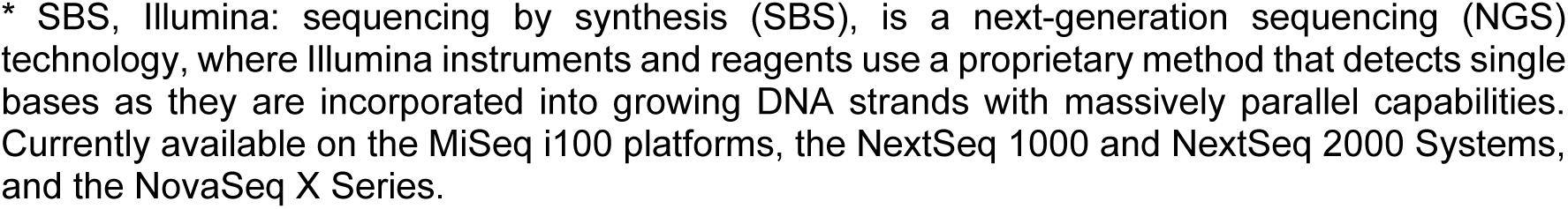
Main results of the studies included in the systematic review.

The main sequencers used in the studies were NovaSeq, Miseq and HiSeq (Illumina), with only one study using Ion Torrent S5 XL (Thermo Fisher Scientific, Waltham, MA, USA) (29).

As for virus families, viral species of medical and veterinary importance identified as part of mosquito viromes were described, belonging to 53 virus families, as well as unclassified virus sequences. After analyzing the 17 studies, vertebrate viruses known to infect mammals naturally were identified, belonging to the Adenoviridae, Anelloviridae, Circoviridae, Coronaviridae, Flaviviridae, Herpesviridae, Papillomaviridae, Reoviridae, Retroviridae, Togaviridae and Rhabdoviridae families (26,39).

The Flaviviridae family was detected in 14 studies, represented by flaviviruses that infect vertebrates and some insect flaviviruses (26). It was observed in the studies that the detection of sequences belonging to the Flaviviridae family is important due to the high number of sequences identified and their phylogenetic proximity to known pathogens, such as St. Louis encephalitis virus (SLEV; Flaviviridae). Louis encephalitis virus (SLEV; Flaviviridae: Flavivirus), West Nile virus (WNV; Flaviviridae: Flavivirus), Zika virus (ZIKV; Flaviviridae: Flavivirus), and dengue virus (DENV; Flaviviridae: Flavivirus).

The studies highlight the importance of research into ISVs (Insect-Specific Viruses) known to be non-pathogenic, which naturally infect insects and replicate exclusively. Although these insect-specific viruses do not appear to replicate in vertebrates, some of them are phylogenetically related to known pathogenic viruses and there is speculation that they may evolve to become new pathogenic viruses in vertebrates (37,40).

Worthy of note are the studies by Calle-Tobón *et al.* (8) who detected in a sample of Aedes aegypti ISVs from the Totiviridae family called the Aedes aegypti toti-like virus and the Australian Anopheles totivirus, and an ISV from the Iflaviridae family called the iflavirus. The studies by Oguzie *et a*l. (34) detected the Tesano Aedes virus (TeAV), a member of the Iflaviridae family, in mosquito pools. As well as the studies by Kubacki *et al.* (30) which detected members of the Dicistroviridae family.

The abundance of viral readings varied greatly according to the virus classification and mosquito species. In the studies, species from viral families known to be associated with plants were detected, among them, the Virgaviridae, Sobemoviridae families (39), the Tombusviridae family (29), the Tymovirdae family (30) and members of the Luteoviridae family (32)..

Viral sequences belonging to the Mimiviridae and Totiviridae families, known to infect protozoa, were also detected (26). In most studies, unclassified viruses represented a considerable number of viral sequences, indicating the need to study them in order to understand their genetic, ecological and evolutionary diversity and their ability to promote future outbreaks (36).

### Quality of evidence

Information regarding the quality of evidence/risk of bias was presented in Table 3 based on the SYRCLE tool for animal studies (20) and adapted according to the guidelines of this tool, which recommends that researchers and collaborators who are going to assess the risk of bias of the included studies discuss in advance and adapt it to the specific needs of their review (20).

**Table 3:** Evaluation of the quality of the studies according to the SYRCLE scale.

| Author / year |  | Selection bias |  |  | Performance Bias |  | Detection Bias |  | Friction Bias | Reporting Bias | Other sources of bias |
| --- | --- | --- | --- | --- | --- | --- | --- | --- | --- | --- | --- |
|  |  | 1 | 2 | 3 | 4 | 5 | 6 | 7 | 8 | 9 | 10 |
| 1. | Sanborn *et al*, 2021 (39). | Y | Y | N | N | N | Y | Y | ? | Y | Y |
|  | Williams *et al*, 2020 (21). | Y | Y | N | N | N | Y | Y | ? | Y | Y |
| 2. | Calle-Tobón *et al*, 2022 (8). | Y | Y | N | N | N | Y | Y | ? | Y | Y |
| 3. | Reuter *et al*, 2016 (28). | Y | Y | N | N | N | Y | Y | ? | Y | Y |
| 4. | Öhlund *et al*, 2019 (29). | Y | Y | N | N | N | Y | Y | ? | Y | Y |
| 5. | Kubacki *et al*, 2020 (30). | Y | Y | N | N | N | Y | Y | ? | Y | Y |
| 6. | He *et al*, 2021 (3). | Y | Y | N | N | N | Y | Y | ? | Y | Y |
| 7. | Thannesberger *et al*, 2021 (31). | Y | Y | N | N | N | Y | Y | ? | Y | Y |
| 8. | Omuoyo *et al*, 2023 (22). | Y | Y | N | N | N | Y | Y | ? | Y | Y |
| 9. | Sadeghi *et al*, 2018 (32). | Y | Y | N | N | N | Y | Y | ? | Y | Y |
| 10. | Ribeiro *et al*, 2019 (33). | Y | Y | N | N | N | Y | Y | ? | Y | Y |
| 11. | Oguzie *et al*, 2022 (34). | Y | Y | N | N | N | Y | Y | ? | Y | Y |
| 12. | Atoni *et al*, 2018 (26). | Y | Y | N | N | N | Y | Y | ? | Y | Y |
| 13. | Faizah *et al*, 2020 (35). | Y | Y | N | N | N | Y | Y | ? | Y | Y |
| 14. | Hameed *et al*, 2021 (36). | Y | Y | N | N | N | Y | Y | ? | Y | Y |
| 15. | Sanborn *et al*, 2019 (37). | Y | Y | N | N | N | Y | Y | ? | Y | Y |
| 16. | Shahhosseini *et al*, 2017 (38) | Y | Y | N | N | N | Y | Y | ? | Y | Y |

Y - YES (low risk of bias); N - no (high risk of bias); ? - uncertain (uncertain risk of bias)

All the studies included in this systematic review were primary, experimental, cross-sectional and prospective. Although this systematic review had limitations in terms of assessing methodological quality, all the studies were classified as having a low risk of bias.

Regarding the selection bias presented in columns 1, 2 and 3 of table 3, it refers to the allocation sequence: In all studies, mosquitoes were identified by species and allocated into groups (pools), which can be seen in table 1. Regarding the basic characteristics: In all the articles, the groups were standardized in terms of the number of specimens, stored in refrigeration to preserve the viral DNA or RNA, and then processed for metagenomic sequencing.

Item 3 presented a high risk of bias, as there was no concealment in the allocation of groups, as it is essential to identify the mosquitoes down to the species, to understand the diversity and ecology of the viruses that will be found after genomic sequencing.

Regarding performance bias, columns 4 and 5 respectively referred to random housing and blinding. In all articles, the groups were distributed by species and stored in refrigeration, without the need for random housing. Regarding blinding, it is important that the researcher has knowledge about the mosquito species; there was no intervention. Therefore, they presented a high risk of bias.

When reporting on detection bias, column 6 deals with the random assessment of the outcome, carried out randomly according to the detection of viruses in the samples. Column 7 shows that blinding was the application of a metagenomic technique in order to detect all the viral sequences present in the sample. Both items presented a low risk of bias.

Regarding attrition bias, in item 8, referring to the incomplete outcome result: No article mentioned the exclusion of mosquito groups in the outcome assessment, we classified the risk of bias as uncertain. With regard to reporting bias, presented in item 9, called selective outcome reporting, the presence of viruses was detected in all the studies. As for other sources of bias, item 10, no article presented other sources of bias. Thus presenting a low risk of bias.

## Discussion

In order to seek evidence of the effectiveness of the viral metagenomics technique in detecting arboviruses in mosquitoes, this systematic review identified 17 articles that obtained promising results in detecting arboviruses in mosquitoes, in addition to the search strategies leading to knowledge of the diversity of virus species in more than 53 viral families.

The eligible studies that make up this systematic review used the viral metagenomics technique as an important tool from the perspective of pathogen surveillance and control (41,42), proving to be efficient in identifying arboviruses, due to the sensitivity in detecting all viral genomes that coexist in a given sample (14), in addition to playing a relevant role in understanding viral diversity and the ecological function that viruses develop in the environment (3,39).

Arboviruses (Arthropod-borne viruses) are viruses transmitted in nature by hematophagous arthropods, with emphasis on dipterans of the Cullicidae family, often maintained in complex cycles involving vertebrates, such as mammals or birds and other vectors that feed on blood. Both humans and animals are infected, and may present diseases ranging from subclinical to fevers, encephalitis, hemorrhagic diseases, with a significant proportion of fatalities (43).

RNA viruses are directly related to the cause of new emerging diseases, as the mutation rate is predominantly high, allowing adaptation to new hosts (30), increasing the transmissibility of viruses by several species of mosquitoes, directly contributing to their geographic distribution (3). In the last decade, deep sequencing technologies have further enabled the discovery of newly identified RNA viruses associated with arthropods, including those discovered in mosquitoes (38).

Thus, the richness and abundance of Culicidae found in a given area, especially those with a history of arbovirus outbreaks, with environments conducive to mosquito breeding, constitute an important reservoir for many different viruses, contributing to the spread and evolution of the virus (3). Mirroring the virome of Culicidae populations through metagenomic analysis aims to explore the spatial and temporal dynamics of virus diversity in mosquitoes, in order to understand virus-virus interactions and identify new strategies for preventing arbovirus diseases. (33).

Therefore, the mosquito virome arouses interest due to the possibility of detecting new viruses, which allows a critical look at the ecology of emerging and reemerging pathogens that circulate in vector populations (21). In particular, the vectors of human viruses are relevant for identifying potential areas of disease outbreaks and for understanding the dynamism of mosquitoes in the genetic evolution of viruses, offering observable ecological correlates, attracting permanent surveillance attention through the viral metagenomics of mosquitoes (30,39).

The worldwide distribution and population density of Culicidae favor viral transmission and increase the risk of viral diseases. Among the various species shown in Table 1, *Ae. albopictus* and *Ae. aegypti* are vectors of several clinically important viruses, such as ZIKAV, DENV, CHIKV and West Nile virus (WNV) (30).

In the studies by Calle-Tobón *et al*. (8) in Medellín, Colombia, 21 types of viruses related to ISVs, arboviruses and plant viruses belonging to nine different viral families were detected through metagenomics in the species of *Ae. aegypti* and *Ae. albopictus*: Flaviviridae, Totiviridae, Reoviridae, Phenuiviridae, Iflaviridae, Bromoviridae, Metaviridae, Xinmoviridae and Orthomyxoviridae; however, six viruses remained unclassified. Only one *Ae. aegypti* sample was positive for DENV3.

According to studies by Atoni *et a*l. (26), carried out in Kenya and southwest China, 240 pools were grouped predominantly of *Culex quinquefasciatus* and *Culex tritaeniorhynchus* species. After genome sequencing, followed by bioinformatics analysis, a total of 30 virus families and a general group of unclassified viruses were detected. The vertebrate viruses known to naturally infect mammals and replicate in mammalian cell lines are highlighted, including Adenoviridae, Anelloviridae, Circoviridae, Coronaviridae, Flaviviridae, Herpesviridae, Papillomaviridae, Reoviridae, Retroviridae, Togaviridae and Rhabdoviridae. Dengue Fever 2 virus was detected in a sample in China and later confirmed by PCR.

Studies by Thannesberger *et al*. (31) focused on mosquito collection in Barbados during the peak of the 2016 ZIKV epidemic, using viral metagenomics in *Ae. aegypt*i mosquitoes, resulting in the detection of two distinct ZIKV genotypes.

Among other eligible studies, Faizah *et al*. (35) collected mosquitoes in Japan in order to isolate the Japanese encephalitis virus, processing samples from *C. tritaeniorhynchus, C. pseudovishnui, C. vishnui* and *C. inatomii* mosquitoes. The metagenomic analyses comprised double-stranded RNA (dsRNA) viruses belonging to the families Totiviridae, Partitiviridae and Chrysoviridae; positive-sense (+) ssRNA single-stranded RNA viruses belonging to the families Flaviviridae and Iflaviridae and unclassified groups; and negative-sense (−) ssRNA viruses from the families Xinmoviridae and Rhabdoviridae and unclassified groups.

Other studies that showed positive results for JEV were conducted in the Yunnan–Myanmar border region of China, in which the species *Cx. Tritaeniorhyncus, Ae. aegypti, An. sinensis* and *Ar. subalbatus* were subjected to NGS (36). Thus, as well as China, in the Shaanxi-Gansu-Ningxia region, species *Culex pipiens, Culex tritaeniorhynchus, Anopheles sinensis,* and *Aedes sp* were collected (3). And NGS of 260 pools of mosquitoes of the species *Cx tritaeniorhynchus* and *Cx bitaeniorhynchus* collected at Camp Humphreys, Republic of Korea (39).

In this sense, technological advances in next-generation sequencing (NGS) have provided an expansion of understanding of the richness, abundance, viral evolution and dynamics of pathogens present in mosquitoes (3,8,21), without the need for laboratory culture and viral isolation, making it possible to identify any viruses present in a given sample, which makes it an important tool in the discovery and surveillance of viruses of medical and veterinary importance (3,36,37,44).

With NGS, the nucleic acids present in a given sample are sequenced, followed by analysis in bioinformatics in order to identify viral sequences by alignment with reference genomic databases. Thus, sequencing platforms are sensitive in detecting arboviruses, providing information on genetic evolution or sequence similarity in relation to viral marker genes (31).

When discussing the application of metagenomics in the detection and characterization of new viruses, the studies by Shahhosseini *et al.* (38) in a pool composed of 25 female specimens of *Aedes cantans*, a mosquito species known to be involved in the transmission of several arboviruses in Europe, were captured in the city of Hamburg, Germany, and were positive for rhabdovirus after being subjected to metagenomic analysis. The virus was genetically related to recently discovered mosquito-associated rhabdoviruses. The studies by Reuter *et al.* (28) also detected three pools of mosquitoes positive for a new rhabdovirus that contained only *Ochlerotatus sp*. mosquitoes collected in Hungary, which were detected and genetically characterized.

Among the results presented in Table 2, the viruses belonging to the genera *Alphavirus* of the Togaviridae family and *Flavivirus* of the Flaviviridae family stand out; and other arboviruses of clinical importance belonging to the families Bunyaviridae, Reoviridae and Rhabdoviridae’ (5).

Faced with this diversity of constantly mutating virus populations, metagenomics enables the comparison of virulence characteristics with available genomic databases, determining susceptibility to diseases and enabling the control and tracking of diseases, and the possibility of predicting and preventing future threats to public health (3).

In addition to arboviruses, another point discussed in the studies that make up this systematic review is related to insect-specific viruses (ISVs). With the improvement of sequencing platforms, the investigation and discovery of insect-specific viruses has been significantly expanded through metagenomics. Mosquitoes have been the main focus of ISV screening (45).

ISVs are restricted to arthropods and are unable to replicate in vertebrate cells or tissues (22). The prevalence of ISVs in wild mosquito populations is underscored by their ability to suppress, increase or have no effect on the replication of medically important arboviruses, potentially affecting vector competence, contributing to their use as biocontrol agents (44).

Although the mechanism by which ISVs modulate arboviruses is unclear, no ISV is known to prevent transmission of all common arboviruses of veterinary and human importance (22,32,41). In this sense, many ISVs are shown to be genetically related to the phylogeny of pathogenic viruses. With this close genetic similarity, it is possible to clarify the potential for interference in the transmission of pathogens or the dynamics of replication in the vector, where similar viruses can block each other through competition (44,45).

Thus, we envision studies that could lead to advances in the understanding of ISVs related to their effects on arbovirus infection and transmission, and in the use of mechanisms in the biological control of vectors, as well as providing the emergence of new vaccine platforms (32,41).

Among the diversity of viruses detected in the mosquito virome, the studies highlight the abundant detection of unclassified viruses (37), made visible by recent advances in random amplification technologies, and metagenomic surveillance has enabled the expansion in the number of new, and often unclassified, viruses (38).

NGS advances have enabled the molecular and taxonomic identification of viruses, assigned by the International Committee on Taxonomy of Viruses (ICTV). However, the results of the comparison of the viral content of the contigs are classified in a general group of unclassified viruses, due to the need for studies that show coherent information from the perspective of their ecology and morphological composition (26).

There is little information available on unclassified viruses and their evolution in mosquitoes. The discovery and characterization of these viruses highlights the need for constant vigilance in preventing future outbreaks, due to the potential for mutation, which contributes to the emergence of new diseases in vertebrates (36).

Among the main challenges pointed out in metagenomic studies is the limited sequence information in the databases for viral families and genera (8). Thus, the lack of reference sequences for most mosquito species is also a factor against the computational removal of RNA readings (42).

Another problem raised in the studies refers to the analysis of metagenomics of entire mosquitoes, due to the size of viruses that can be included in the analysis, such as plant viruses belonging to their diet and viruses from parasitic or commensal organisms that reside inside or under the mosquito (29).

## Conclusion

In view of what was proposed in this systematic review, the efficacy of the viral metagenomics technique in the detection of arboviruses was demonstrated in the 17 eligible studies that make up this review, both in the detection of pathogens and in the characterization of viral diversity, with the aim of understanding the ecology or epidemiology of these viruses. Metagenomics has emerged as an important tool responsible for expanding the taxonomic identification of viruses based on studies that seek to understand viral genomes in recent years.

Metagenomics has simplified the detection of viruses in mosquito samples, enabling constant epidemiological surveillance, predicting and preventing outbreaks, due to its sensitivity in detecting viral sequences without the need for a large sample and isolation by culture. On the other hand, the high financial costs of acquiring and maintaining NGS technology restrict its use to a few research laboratories.

Despite the few studies on a global scale, there is a need to deepen our knowledge of the mosquito virome in order to better meet the social, economic and health needs of globalization.

## Acknowledgement

We thank CAPES (Coordination for the Improvement of Higher Education Personnel) for granting the scholarship (#88887.823555/2023-00). Ao Departamento de Extensão da Universidade Federal do Ampá (DEX-UNIFAP). Dr. Shirley Vasconcelos Komninakis and her research group at the Retrovirology Laboratory at the Federal University of São Paulo for their academic support in conducting and supervising this work. And to Dr. Raimundo Nonato Picanço Souto, from the laboratory of Arthropoda, Federal University of Amapá, for his guidance and supervision of this important study.

## Conflicts of Interest

The Authors declare no competing interests

## Notes

### Competing Interest Statement

The authors have declared no competing interest.

